# A HISTORY OF ADOLESCENT OXYCODONE SELF-ADMINISTRATION: NEURAL MECHANISMS UNDERLYING NEUROCOGNITIVE IMPAIRMENTS

**DOI:** 10.64898/2026.01.19.700403

**Authors:** Kristen A. McLaurin, Taylor R. Elder, Hailong Li, Brendan J. Veenstra, Kyler J. Nelson, Charles F. Mactutus, Jill R. Turner, Rosemarie M. Booze

## Abstract

Maturation of the prefrontal cortex and higher-order cognitive processes is ongoing through adolescence and young adulthood; developmental periods associated with the nonmedical use of prescription opioids. Initiation of oxycodone (OXY) self-administration during adolescence induces long-term dose-dependent impairments in neurobehavioral and neurocognitive development. To date, however, the neural mechanisms underlying neurocognitive impairments induced by adolescent prescription opioid use disorder (APOUD) have not been systematically evaluated. A DiOlistic labeling technique, in concert with sophisticated neuronal reconstruction software, revealed a prominent sex- and dose-dependent population shift in the morphological parameters of dendritic spines in pyramidal neurons in the anterior cingulate cortex (ACC) of rodents with a history of adolescent OXY self-administration relative to their control counterparts. Similarly, the biochemical mechanisms underlying dendritic spine dysmorphology, which were investigated using quantitative polymerase chain reaction, were dependent upon biological sex. In male rodents, a dose-dependent decrease in mRNA expression of *GRIN1,* a N-methyl-D-aspartate receptor, was observed in animals with a history of adolescent OXY self-administration relative to controls. In contrast, female rats with a history of adolescent OXY self-administration exhibited increased mRNA expression of *cdk5*, a protein kinase involved in the regulation of pre- and post-synaptic function; a transcriptional alteration that may result from excessive glutamate accumulation. Fundamentally, transcriptional dysregulation and dendritic spine dysmorphology mechanistically underlie, at least partly, APOUD-induced neurocognitive impairments. Collectively, results support that sex-dependent glutamatergic dysregulation may underlie dendritic spine dysmorphology in the ACC resulting in APOUD-induced neurocognitive impairments. Glutamatergic dysregulation, therefore, may afford a key target for the development of novel therapeutics.

**SIGNIFICANCE STATEMENT:** Individuals with a history of adolescent prescription opioid use disorder (APOUD) are afflicted with profound neurocognitive impairments. Herein, sex-dependent glutamatergic dysregulation may underlie dendritic spine dysmorphology in the ACC resulting in APOUD-induced neurocognitive impairments, thereby affording a key target for the development of novel therapeutics.

## INTRODUCTION

Adolescence (12-17 years of age) and young adulthood (18-25 years of age) are developmental periods of transition from childhood to adulthood characterized by significant maturational changes in the brain; ontogenetic changes that render these individuals uniquely vulnerable to substance use initiation.^1^ Indeed, nonmedical use of prescription opioids peaks during late adolescence.^2–3^ Consequently, in 2023, 2.2% of adolescents and 2.5% of young adults reported misuse of prescription pain relievers (e.g., Codeine, Oxycodone (OXY)), corresponding to an estimated 570,000 and 841,000 individuals, respectively.^4^ The long-term consequences of adolescent prescription opioid use, misuse, and/or abuse cannot be understated, as it is associated with an increased risk of subsequent opioid use disorder (OUD)^5–6^ or substance use disorder,^7^ progression to heroin use,^8–9^ and development of neurocognitive impairments.^10–11^

Indeed, generalized neurocognitive dysfunction is one of the devastating consequences associated with OUD.^12–13^ Cross-sectional studies have provided a wealth of knowledge on neurocognitive impairments in adults with prescription OUD (POUD), characterized by prominent alterations in attention,^14^ memory,^14^ and executive functions.^15–16^ Extrapolating cross-sectional findings to developmental processes, however, is inferentially fraught.^17^ Hence, a longitudinal study was recently undertaken, whereby chronic oral adolescent OXY self-administration selectively disrupted neurobehavioral (i.e., locomotion, startle reactivity) and neurocognitive (i.e., preattentive processes) development.^11^ The repercussions of adolescent POUD (APOUD) are long-lasting, with deficits in preattentive processes,^10–11^ sustained attention,^10–11^ and spatial memory^10,18^ persisting into adulthood. Nevertheless, the neural mechanisms underlying APOUD-induced neurocognitive impairments remain unclear.

The prefrontal cortex (PFC), a brain region involved in higher-order cognitive processes,^19^ undergoes prolonged maturation with continued development through adolescence and young adulthood. Early prenatal development of the PFC is a highly meticulous process characterized by rapid neuronal production, postmitotic neuronal migration, and neuronal differentiation.^20^ Prefrontal synaptogenesis follows thereafter, whereby the dynamic process begins during the prenatal period and continues postnatally, resulting in a surge in synapse formation and exuberant connectivity. Refinement of the PFC during adolescence and young adulthood, in sharp contrast, is characterized by regressive processes, including dendritic pruning^21^ and excitatory synapse elimination;^22–25^ events that aid in the refinement of neural circuits and formation of mature connections.

Dendritic spines, which serve as the postsynaptic compartment of most functional excitatory synapses,^26–27^ are highly dynamic, specialized protrusions emanating from dendrites. Morphologically, dendritic spines are characterized by a long, thin dendritic spine neck and a bulbous dendritic spine head. The morphology of the dendritic spine head, in particular, is tightly coupled to dendritic spine function, whereby a positive correlation between the dendritic spine head and area of the postsynaptic density (PSD) has been reported.^28–29^ Indeed, the PSD, which contains high concentrations of glutamatergic receptors (i.e., N-methyl-D-aspartate receptors (NMDAR), α-Amino-3-hydroxy-5-methyl-4-isoxazolepropionic acid receptors (AMPAR)) and scaffolding proteins (e.g., PSD-95, Homer-1, Shank3), is fundamentally involved in the efficient transduction of neurotransmitter signals into postsynaptic neuron responses. In turn, key scaffolding proteins (e.g., PSD-95^30^, Liprinα1)^31^ are phosphorylated by cyclin-dependent kinase 5 (CDK5). Thus, given the strong structure-function relationship of dendritic spines, examination of continuous dendritic spine morphological parameters may reveal a key neuropathophysiological mechanism underlying persistent neurocognitive impairments induced by APOUD.

Hence, the rationale was built for establishing if, and/or how, oral adolescent OXY self-administration alters the structure and/or biochemical signaling of pyramidal neurons in the anterior cingulate cortex (ACC), a subregion of the medial PFC (mPFC), during adulthood. The structure of pyramidal neurons, and associated dendritic spines, in the ACC were examined using an innovative ballistic labeling technique in concert with sophisticated neuronal reconstruction software. Quantitative polymerase chain reaction (qPCR) techniques were also implemented to evaluate alterations in synaptic scaffolding proteins (i.e., SAP-102: *dlg3,* PSD-95: *dlg4*), a protein kinase localized to postsynaptic compartments (i.e., cdk5)^32–33^ and glutamatergic receptor gene expression. Statistical analyses were conducted to address two fundamental questions, including 1) *Does a history of oral adolescent OXY self-administration alter the dependent variable of interest?* and 2) *Are changes in the dependent variable of interest dependent upon the dose of OXY self-administered by the rodent?* Elucidating the neural mechanisms underlying APOUD-induced neurocognitive impairments is vital for the development of novel therapeutics.

## MATERIALS AND METHODS

### Experimental Design

An experimental design timeline, including oral OXY self-administration, as well as neurocognitive and neuroanatomical evaluations, is illustrated in Figure 1A. A longitudinal evaluation of sensorimotor gating, indexed using prepulse inhibition (PPI), and locomotor activity, indexed using open field activity, which was utilized for regression analyses, was previously published.^11^

**Figure 1.**
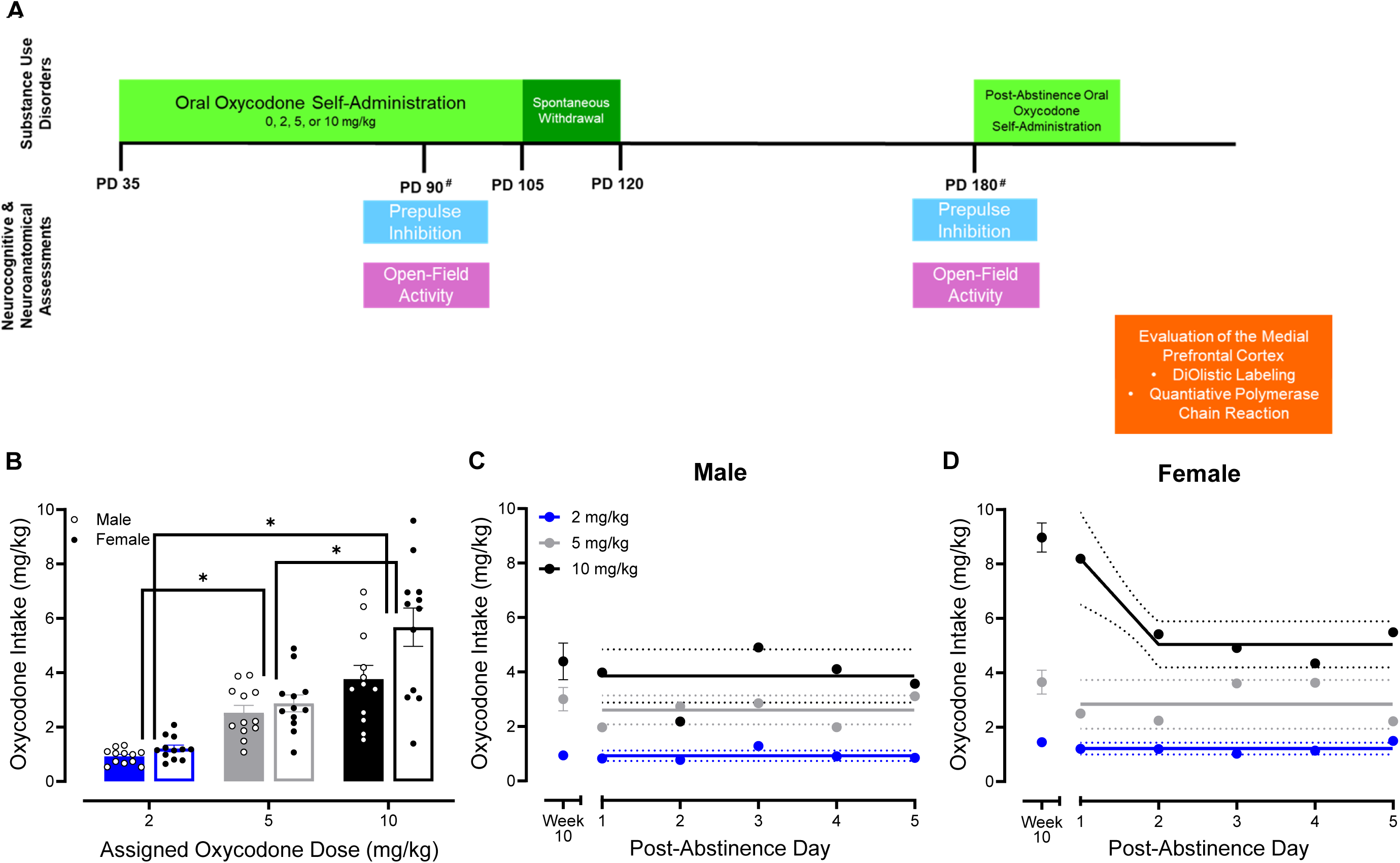
Experimental Timeline and Post-Abstinence Oxycodone (OXY) Intake. (**A**) A schematic of the experimental design. Adolescent oral OXY self-administration and longitudinal neurocognitive assessments, indicted by asterisks (#), were previously published by McLaurin et al.^11^ Assessments of prepulse inhibition and open-field activity at postnatal day (PD) 90 and PD 180 were reanalyzed in a novel manner (i.e., regression analyses). (**B**) Average (Mean (X) ± Standard Error of the Mean) OXY intake (mg/kg) during the five-day post-abstinence period is illustrated as a function of assigned dose group (i.e., 2, 5, or 10 mg/kg) and biological sex (i.e., Male or Female). (**C,D**) OXY consumption (mg/kg; Mean (X) ± 95% Confidence Intervals (CI)) during the post-abstinence period is shown as a function of day and biological sex. Intake during Week 10 of the adolescent period, previously published in McLaurin et al.,^11^ was not included in the analysis, but is included as a reference. Sample sizes for OXY intake are as follows: 2 mg/kg, *n*=24 (male, *n*=12, female, *n*=12), 5 mg/kg, *n*=24 (male, *n*=12, female, *n*=12), and 10 mg/kg, *n*=24 (male, *n*=12, female, *n*=12). OXY intake CIs are illustrated with dotted lines; non-overlapping 95% CIs indicate statistical significance. \**p* ≤ 0.05.

### Animals

Fischer F344/N rats, acquired from Envigo Laboratories Incorporated (Currently Inotiv; Indianapolis, IN) and housed with their biological dam (*N*=15 Litters), arrived at the animal vivarium between postnatal day (PD) 7 and PD 9. At approximately PD 21, rodents were weaned and pair- or group-housed with animals of the same sex for the duration of the experiment. To preclude violation of the assumption of independence, the goal was to select one rat per sex for each OXY dose from each litter (i.e., four male rats per litter and four female rats per litter). A timed water restriction schedule, whereby animals were limited to fluid access for approximately eight hours per day, was implemented at approximately PD 32. Rats had *ad libitum* access to rodent food (Pro-Lab Rat, Mouse, Hamster Chow #3000) for the duration of the study.

The University of South Carolina Institutional Animal Care and Use Committee approved the experimental procedures under federal assurance (#D16-00028). Rodents were maintained in AAALAC-accredited facilities in accordance with recommendations in the Guide for the Care and Use of Laboratory Animals of the National Institutes of Health. Targeted environmental conditions for the animal colony were 21°± 2°C, 50% ± 10% relative humidity, and a 12-h light: 12-h dark cycle (Lights on at 0700 h EST).

### Oral Oxycodone Self-Administration

Baseline neurobehavioral (i.e., locomotor activity) and neurocognitive (i.e., PPI) assessments at PD 30 were conducted to ensure unbiased treatment assignment.^11^ Rodents were subsequently assigned to one of four OXY self-administration dose groups (i.e., 0, 2, 5, or 10 mg/kg) yielding sample sizes of 0 mg/kg, *n*=33 (male, *n*=17, female, *n*=16), 2 mg/kg, *n*=24 (male, *n*=12, female, *n*=12), 5 mg/kg, *n*=24 (male, *n*=12, female, *n*=12), and 10 mg/kg, *n*=24 (male, *n*=12, female, *n*=12).

During the two-bottle choice experimental paradigm, which was adapted from Zanni et al.^34^, rodents had free access to water and/or OXY during the light cycle for approximately eight hours per day. Rodents were singly housed during the eight-hour oral OXY self-administration session and were pair- or group-housed during the 16-hour fluid restriction period. To determine liquid intake, each water bottle was weighed prior to, and upon completion of, free access (i.e., Bottle Weight (g) A.M. – Bottle Weight (g) P.M.).

Three days before beginning oral OXY self-administration, male and female rats were habituated to the two-bottle choice experimental paradigm. During habituation, rodents had access to water in a left and right bottle in a hopper with food placed in between.

At approximately PD 35, an age in rats that corresponds to the adolescent and young adult period in humans,^35^ rodents began oral OXY self-administration. Under two-bottle choice conditions, male and female F344/N rodents had free access to water and OXY dissolved in water (0, 2, 5, or 10 mg/kg) for ten weeks (i.e., through PD 105); the OXY was generously gifted by the National Institute on Drug Abuse Drug Supply Program (Rockville, MD). Two parameters, including the average liquid intake during the habituation period (i.e., 15 mL) and the animal’s biweekly body weight measurement, were used to determine the OXY concentration in each bottle. Animals titrated their OXY intake with a goal of individual rodent intake of approximately 2, 5, or 10 mg/kg/day if the rat consumed the entire 15 mL from the OXY bottle. For half of the experimental animals, the OXY bottle was placed in the left position for Weeks 1 to 5 and in the right position for Weeks 6 to 10; bottle positions were reversed in the other half of the experimental animals. During the adolescent oral OXY self-administration period, both male and female animals voluntarily self-administered OXY, whereby the experimental paradigm induced significant escalation of OXY intake across time.^11^

After a period of abstinence (i.e., at least 10 weeks), rodents received free access to water and OXY dissolved in water (0, 2, 5 or 10 mg/kg) under a two-bottle choice experimental paradigm for five days. With regards to OXY dose, rats were provided with voluntary access to the same dose they were assigned at PD 35. Bottle positions were reversed daily. Rodents were humanely euthanized at least one hour after receiving free access to water and OXY on the sixth day.

### Post-Mortem Analyses of the Prefrontal Cortex

To elucidate the neural mechanisms underlying neurocognitive dysfunction induced by OXY relapse, two neuroanatomical assessments, including ballistic labeling and quantitative polymerase chain reaction, were undertaken. Average OXY intake during the tenth week of adolescent OXY self-administration was utilized to select a subset of animals for each neuroanatomical assessment with the goal of representing the spectrum of OXY intake observed.

### Ballistic Labeling

#### Estrous Cycle Tracking

Immediately prior to euthanasia, cellular cytology in vaginal smears was evaluated using a 10× light microscope. The goal was to sacrifice female rodents during the diestrus phase of the estrous cycle to decrease potential hormonal variability.

#### Tissue Preparation

After deeply anesthetizing animals with sevoflurane (5%; Abbot Laboratories, North Chicago, IL, USA), rodents were transcardially perfused (Control, *n* = 12 (male, *n* = 6, female, *n* = 6), 2 mg/kg, *n =* 12 (male, *n* = 6, female, *n* = 6), 5 mg/kg, *n* = 12 (male, *n* = 6, female, *n* = 6), and 10 mg/kg, *n* = 12 (male, *n* = 6, female, *n* = 6)). The rat brain was removed, post-fixed in 4% paraformaldehyde for 10 minutes, and sliced coronally (500 μm) using a rat brain matrix (ASI Instruments, Warren, MI, USA). Coronal brain slices were placed into a 24-well cell culture plate (Corning, Tewksbury, MA, USA) with 1mM phosphate-buffered saline (PBS). PBS was removed from the targeted wells before ballistic labeling.

#### Ballistic Labeling

The structure of pyramidal neurons, and their associated dendritic spines, in the ACC, a subregion of the mPFC, were visualized using ballistic labeling as detailed in Li et al.^36^ In brief, Tefzel tubing was coated with a polyvinylpyrrolidone solution (i.e., 10 mg of polyvinylpyrrolidone per 1 mL ddH_2_O) for 20 minutes; the PVP solution was subsequently expelled from the Tefzel tubing. To prepare ballistic cartridges, 170 mg of tungsten beads (Bio-Rad, Hercules, CA, USA) and 6 mg of DiIC19(3) dye (Invitrogen, Carlsbad, CA, USA) were independently dissolved in 250 and 300 μL of 99.5% methylene chloride (Sigma-Aldrich, St. Louis, MO, USA), respectively. The tungsten bead suspension was pipetted onto a glass slide and air dried. The DiIC19(3) dye mixture was subsequently pipetted on top of the dried tungsten bead suspension; the two layers were mixed thoroughly, allowed to dry, and divided into two 1.5 mL centrifuge tubes filled with ddH_2_O. The tungsten bead/DiIC19(3) dye mixture was homogenized using sonication, slowly drawn into the polyvinylpyrrolidone-coated Tefzel tubing, and placed into the tubing preparation station (Bio-Rad). After rotating for one minute, all water was removed from the polyvinylpyrrolidone-coated Tefzel tubing; the tubing was rotated for an additional 30 minutes under nitrogen gas (0.5 L per minute). Ballistic cartridges were cut into 13 mm lengths and stored in the dark.

At the time of experimentation, prepared ballistic cartridges were loaded into the Helios gene gun (Bio-Rad). The applicator of the Helios gene gun, which had a piece of filter paper between two mesh screens, was placed vertically at the center of the targeted tissue well and approximately 1.5 cm away from the brain tissue. A high-velocity stream of helium (Output Pressure: 90 psi) was utilized to inject the DiIC19(3) dye-coated tungsten beads into brain tissue; brain tissue slices located approximately 3.7 mm to 2.2 mm anterior to Bregma^37^ were labeled. The labeled brain tissue was washed three times with 100 mM PBS, stored at 4°C in the dark for three hours, and mounted (Pro-Long Gold Antifade; Invitrogen) onto a glass slide.

#### Confocal Imaging

Pyramidal neurons in the ACC were imaged using a Nikon TE-2000E confocal microscopy system outfitted with Nikon’s EZ-C1 software (version 3.81b). Imaging was restricted to the ACC subregion of the rat mPFC, whereby most pyramidal neuron images were obtained from a slice approximately 2.52 mm anterior to Bregma.^37^ Specifically, z-stack images of neurons were obtained by a blinded experimenter using a 60× oil objective with a numerical aperture of 1.4 and a z-plane interval of 0.15 μm (Figure 2).

**Figure 2.**
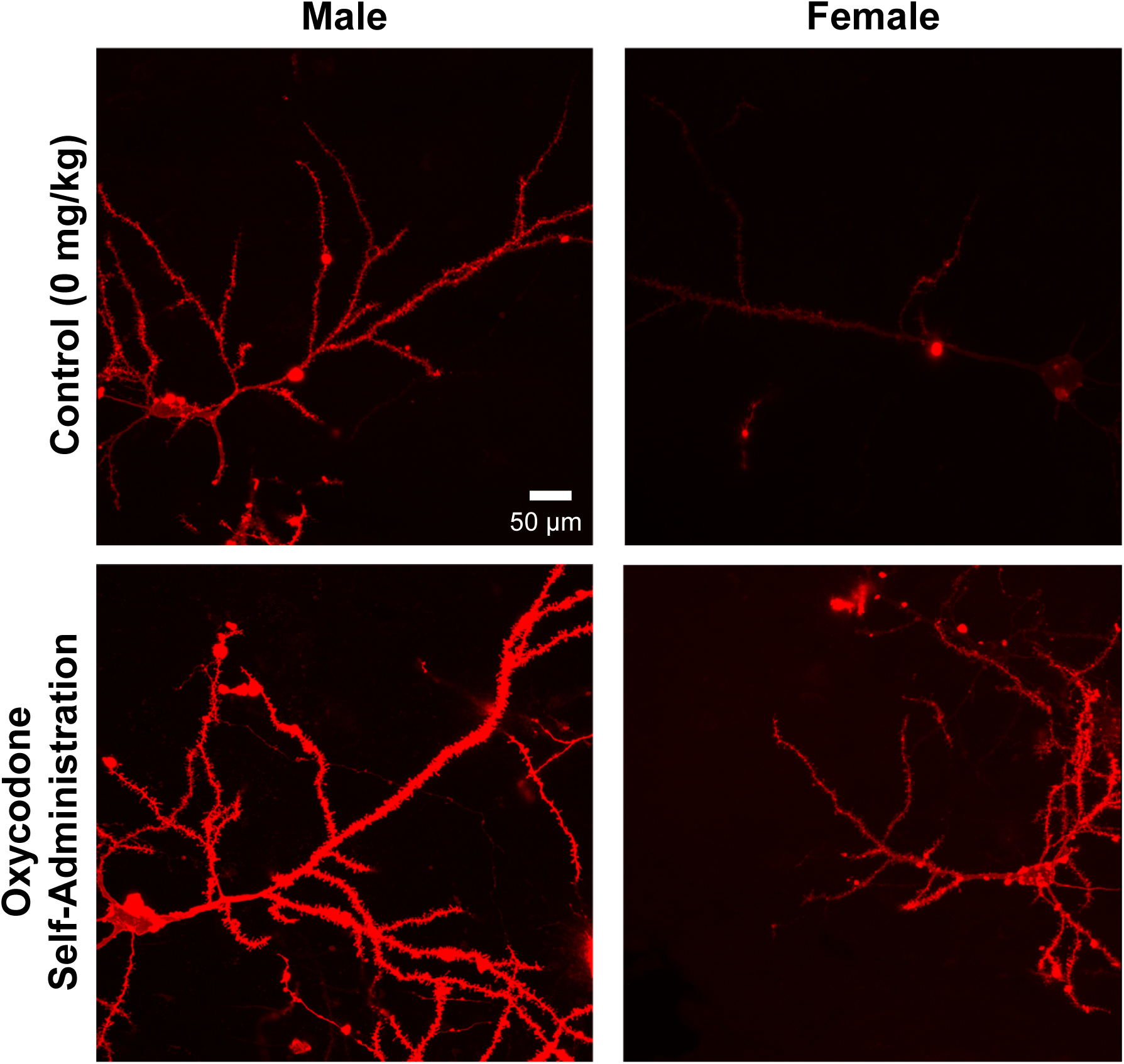
DiOlistic labeling of pyramidal neurons in the anterior cingulate cortex (ACC), a subregion of the medial prefrontal cortex. Pyramidal neurons in male (**A,C**) and female (**B,D**) rodents were characterized by one apical dendrite and multiple basilar dendrites. Female rodents with a history of adolescent oxycodone self-administration exhibited decreased neuronal arbor complexity relative to their control counterparts.

#### Neuronal and Dendritic Spine Analysis

An independent, blinded investigator utilized sophisticated neuronal reconstruction software (Neurolucida 360®; MicroBrightfield, Williston, VT, USA) to analyze confocal images of pyramidal neurons and associated dendritic spines from the ACC. After blinding, stringent selection criteria (e.g., continuous dendritic staining, low background/dye clusters) defined by Li et al.^36^ were used to select one neuron from each animal, yielding the following sample sizes: Control, *n* = 9 (male, *n* = 5, female, *n* = 4), 2 mg/kg, *n =* 8 (male, *n* = 5, female, *n* = 3), 5 mg/kg, *n*=9 (male, *n* = 5, female, *n* = 4), and 10 mg/kg, *n* = 11 (male, *n* = 6, female, *n* = 5).

The morphological parameters of pyramidal neurons in the ACC were established using the classical Sholl analysis,^38^ whereby the number of times dendrites intersect with successive concentric spheres placed around the cell body was counted. Boundary conditions for dendritic spines were established based on previously published literature (volume, 0.05 to 0.85 µm;^39^ backbone length, 0.4 to 4.0 µm;^29,40^ head diameter, 0 to 1.2 µm);^41^ dendritic spines failing to meet any of the boundary conditions were excluded from the analysis. Two parameters, including dendritic spine backbone length (μm) and head diameter (μm), were utilized to evaluate dendritic spine morphology.

### Quantitative Polymerase Chain Reaction

#### Tissue Preparation

Rodents were deeply anesthetized using 5% sevoflurane (Abbot Laboratories, North Chicago, IL, USA) and rapidly decapitated (Control, *n* = 13 (male, *n* = 8, female, *n* = 5), 2 mg/kg, *n =* 12 (male, *n* = 6, female, *n* = 6), 5 mg/kg, *n* = 12 (male, *n* = 6, female, *n* = 6), and 10 mg/kg, *n* = 11 (male, *n* = 5, female, *n* = 6)). The rat brain was removed and frozen in liquid nitrogen within five minutes of sacrifice.

#### RNA Isolation and cDNA Synthesis

Rodent tissue from the PFC was dissected, weighed, and homogenized in TRIzol^®^ (Invitrogen, Waltham, MA; 1 mL TRIzol^®^ per 50-100 mg of tissue). RNA was isolated from the homogenized tissue using the Invitrogen PureLink RNA Mini Kit (Invitrogen) according to the manufacturer’s instructions; the yield of the purified RNA was determined using a nanodrop (NanoDrop One/One Microvolume UV-Vis Spectrophotometer, Thermo Scientific, Waltham, MA). Complementary DNA was subsequently synthesized from 500 ng of the extracted RNA using Oligo(dT) primer (Promega, Madison, WI) and SuperScript II Reverse Transcriptase (Invitrogen).

#### Primer Design

Oligonucleotide primer sequences were designed using the National Center for Biotechnology Information Primer-Designing Tool and checked for specificity using the Basic Local Alignment Search Tool database. Primers were obtained from Integrated DNA Technologies (Coralville, IA); the primers evaluated, their respective sequences, and sample sizes for each gene are reported in Table 1.

**Table 1.**
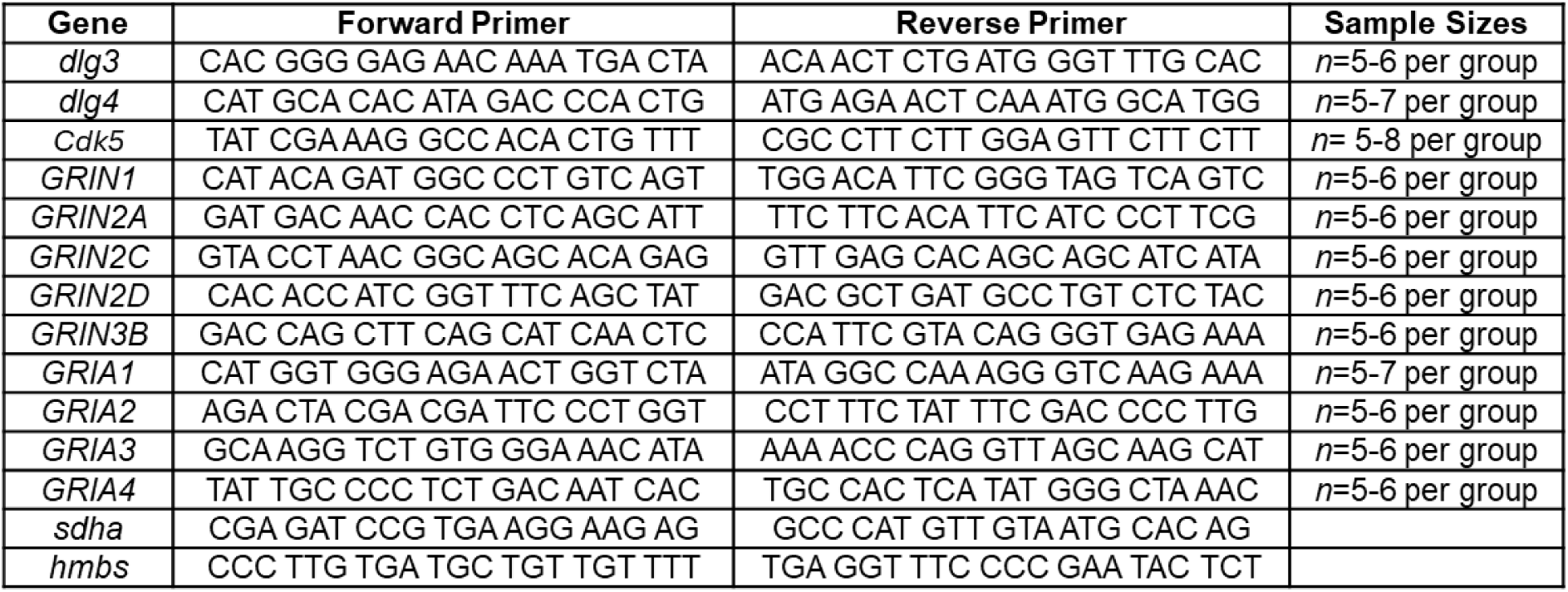
Sequence of primers used in quantitative reverse transcriptase polymerase chain reaction.

#### SYBR-Green QPCR

A primer mix, composed of 100 µL of 20 µm forward primer, 100 µL of 20 µm reverse primer, and 800 µL of nuclease-free water, was prepared for each primer evaluated. Seven microliter qPCR reactions were assembled using 1 µL cDNA, 2 µL of primer mix, and 4 µL of 2x SYBR Green Mix (Applied Biosystems, Waltham, MA) in a 384-well plate; transcript measurements were conducted in triplicate. Real-time PCR reactions were performed with the Bio-Rad CFX384 Real-Time PCR Detection System (Bio-Rad Laboratories, Hercules, CA) using the following cycling conditions: 95°C for 10 minutes, 40 cycles of 95°C for 15 seconds followed by 60°C for 1 minute, and a final melt curve of 60°C to 95°C in 0.5°C increments over 10 seconds. mRNA levels were determined using the 2^-ΔΔCt^ method.^42^

Two housekeeping genes, including Hydroxymethylbilane Synthase (*HMBS*) and Succinate Dehydrogenase Complex Subunit A (*SDHA*), were run on each qPCR plate. The mRNA expression of *HMBS* and *SDHA* did not systematically change across OXY dose and exhibited strong stability, evidenced by low Ct value standard deviations. Thus, target genes were normalized to either *HMBS* or *SDHA* after assessing the Ct value for genes of interest. Gene expression values were normalized to male control (i.e., 0 mg/kg) rodents.

### Statistical Analysis

The neural mechanisms underlying APOUD-induced neurocognitive impairments were evaluated using two complementary approaches. First, statistical analyses were conducted to determine if and/or how a history of oral adolescent OXY self-administration alters the dependent variable of interest. Second, analytic techniques were utilized to determine if and/or how the dose of OXY self-administered by the rodent is associated with the dependent variable of interest.

Analysis of variance (ANOVA) techniques and regression analyses were utilized to evaluate statistical significance at an alpha level of *p* ≤ 0.05 (SPSS Statistics 29, IBM Corp., Somers, NY, USA; SAS/STAT Software 9.4, SAS Institute, Inc., Cary, NC; GraphPad Prism 10 Software, Inc., La Jolla, CA). Figures were created using GraphPad Prism 10 Software.

A two-way ANOVA with Tukey’s HSD post hoc test (SPSS Statistics 29), repeated measures ANOVA (SPSS Statistics 29) and complementary regression analyses (GraphPad Prism 10) were utilized to analyze OXY intake (mg/kg) during the non-abstinent period. Intake during Week 10 of the adolescent period, previously published in McLaurin et al.,^11^ was not included in the analysis, but is included in Figure 1C and 1D as a reference. Censored data, which resulted from a leaky water bottle (Day 1, 2 mg/kg Male, *n*=1; Day 1, 5 mg/kg Female, *n*=2; Day 2, 10 mg/kg Male, *n*=1; Day 3, 2 mg/kg Female, *n*=2), were resolved using the mean series imputation method.

Neuronal and dendritic spine morphology were analyzed using a mixed effects model with multi-level nesting (Covariance Structure: AR(1))^43^ and generalized linear mixed-effects models with a Poisson distribution (Covariance Structure: UN; SAS/STAT Software 9.4, PROC GLIMMIX), respectively. The number of intersections at successive radii, the dependent variable of interest for neuronal morphology, was transformed by squaring the values.

Univariate ANOVA (SPSS Statistics 29) and regression (GraphPad Prism 10) techniques were utilized to evaluate OXY-induced alterations in mRNA expression. The delta CT value was utilized as the dependent variable of interest for the univariate ANOVA. Outliers that were greater than 2.5 standard deviations away from the mean were removed from the analyses and figures (*GRIN1*: 10 mg/kg Male, *n*=1). Statistical analyses reported for the Main Effect of Biological Sex included OXY Access, not OXY Dose, as the independent variable of interest.

Simple (GraphPad Prism 10) and multiple (SPSS Statistics 29) regression analyses were utilized to evaluate the relationship between proposed underlying neural mechanisms (e.g., *GRIN1* and *dlg3* mRNA expression), as well as between dendritic spine dysmorphology and previously published measures of neurocognition (e.g., visual PPI).^11^ The most extreme residual and two outliers were removed from the analysis evaluating the relationship between *GRIN1* mRNA expression and visual PPI in male rodents. Further, one outlier was removed from the analysis evaluating the relationship between mean dendritic spine backbone length and locomotor activity in female animals.

## RESULTS

### Rodents voluntarily consumed oxycodone following a period of abstinence

After a significant period of abstinence (i.e., at least 10 weeks), male and female rats with a history of oral adolescent OXY self-administration were again provided with access to the same dose of OXY (i.e., 2, 5, or 10 mg/kg) and water under a two-bottle choice conditions for five days. Evaluation of the average OXY intake (mg/kg) across the five-day post abstinence period revealed that consumption occurred in a dose-dependent manner (Figure 1B; Main Effect: OXY Dose [*F*(2,66)=42.6, *p*≤ 0.001, η_p_^2^=0.564]), whereby independent of biological sex, rodents with access to 5 mg/kg OXY consumed a higher dose than those with access to 2 mg/kg (*p* ≤ 0.001). In a similar manner, animals with access to 10 mg/kg OXY consumed a higher dose than those with access to either 2 mg/kg (*p* ≤ 0.001) or 5 mg/kg (*p* ≤ 0.001).

Subsequent analyses, which included time as a factor, revealed that OXY consumption (mg/kg) during the post-abstinence period occurred in a time-, dose- and sex-dependent manner (Day x OXY Dose x Sex Interaction [*F*(8,264)=3.5, *p_GG_* ≤ 0.004, η_p_^2^=0.095] with a prominent linear component [*F*(2,66)=5.6, *p* ≤ 0.006, η_p_^2^=0.145]). Complementary analyses, therefore, were conducted independently for male and female rodents.

In male rats, the assigned OXY dose group, but not time (*p* > 0.05), influenced the voluntary consumption of OXY following a period of abstinence (Figure 1C; Main Effect: OXY Dose [*F*(2,33)=18.1, *p* ≤ 0.001, η_p_^2^=0.522]). Regression analyses confirmed these observations, whereby a horizontal line afforded the best-fit function for the 2, 5, and 10 mg/kg doses; albeit statistically significant differences in the mean of the curve were observed [*F*(2,174)= 35.4, *p*≤ 0.001].

In sharp contrast, OXY intake following a period of abstinence occurred in a time- and dose-dependent manner in female rodents (Figure 1D; Day x OXY Dose Interaction [*F*(8,132)=4.0, *p_GG_* ≤ 0.002, η_p_^2^=0.195] with a prominent linear component [*F*(2,33)=10.9, *p* ≤ 0.001, η_p_^2^=0.398]); an interaction that is driven by the female animals assigned to the 10 mg/kg OXY dose group. Indeed, post-abstinence OXY consumption in female animals assigned to the 10 mg/kg OXY dose group was well-described using a segmental linear regression (*R*^2^ = 0.90), evidenced by a decrease in intake from Day 1 to Day 2, followed by a subsequent plateau. A horizontal line, however, afforded the best fit-function for the 2 and 5 mg/kg doses; albeit statistically significant differences in the mean of the curve were observed [*F*(1,114)= 45.8, *p*≤ 0.001].

### Oxycodone self-administration induced structural alterations to pyramidal neurons in the anterior cingulate cortex in a sex-dependent manner

The classic Sholl analysis,^38^ which quantifies the number of times dendrites intersect with concentric spheres at successive radii, affords an opportunity to infer neuronal arbor complexity. Chronic adolescent oral OXY self-administration induced a sex-dependent alteration in the number of intersections at successive radii (Figure 3A-B; OXY Access x Sex x Radii Interaction, [*F*(17, 561)=1.68, *p*≤0.05]). Complementary analyses, which were conducted by sex, revealed the locus of the interaction, whereby female (Figure 3B; OXY Access x Radii Interaction, [*F*(17, 255)=2.2, *p*≤0.005]), but not male (Figure 3A; *p*>0.05), OXY rodents exhibited a prominent decrease in the number of intersections at successive radii relative to female or male control animals, respectively.

**Figure 3.**
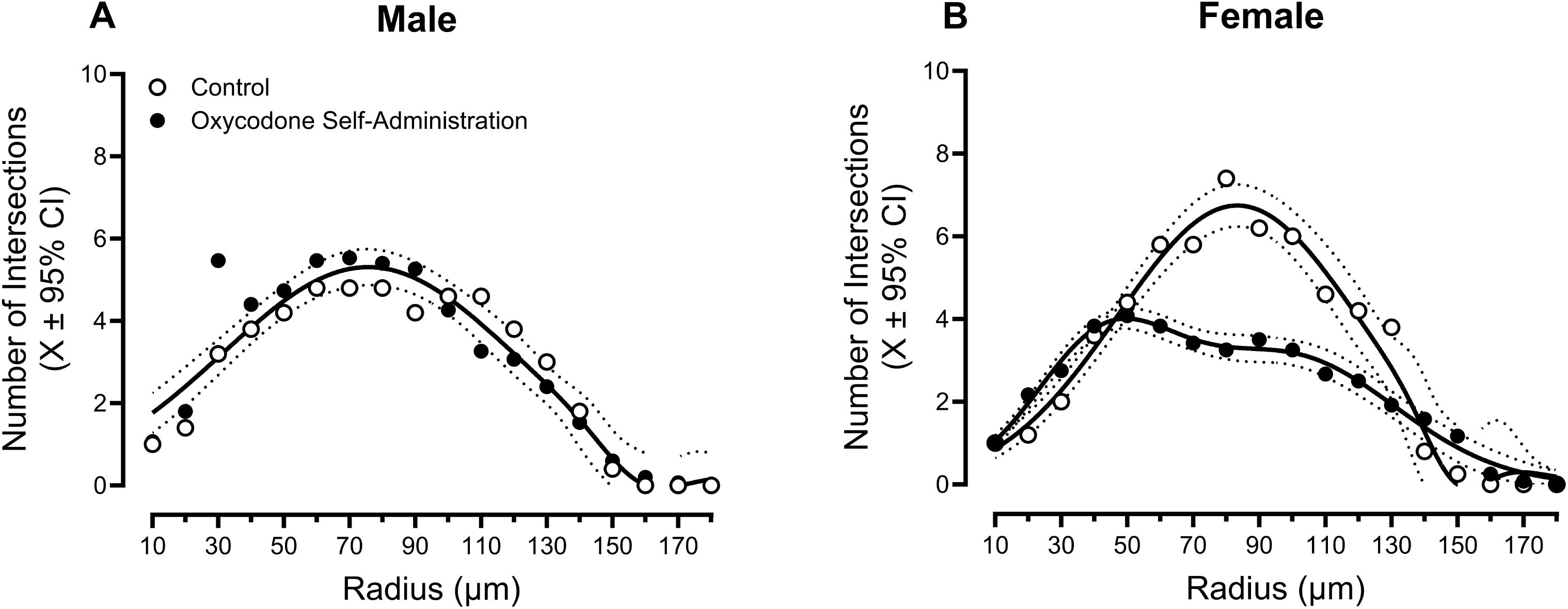
Sholl Analysis. (**A,B**) The number of dendritic intersections occurring every 10 μm from the soma (Mean (X) ± 95% Confidence Intervals (CI)) were measured to evaluate neuronal arbor complexity (Control (i.e., 0 mg/kg): *n*=9 (male, *n*=5, female, *n*=4); Oxycodone Self-Administration: *n*=28 (male, *n*=16, female, *n*=12)). Female (**B**), but not male (**A**), rodents with a history of adolescent oxycodone self-administration exhibited a statistically significant decrease in the number of dendritic intersections relative to their control counterparts evidenced by non-overlapping 95% CIs.

Structural alterations to pyramidal neurons in the ACC were not, however, dose-dependent (Data Not Shown; OXY Dose x Sex x Radii, *p*>0.05). Collectively, a history of chronic adolescent oral OXY self-administration induces significant reductions in neuronal arbor complexity in female, but not male, rodents.

### A sex- and dose-dependent population shift in the morphology of dendritic spines was observed following adolescent oxycodone self-administration

Two parameters, including backbone length (µm) and head diameter (µm), were utilized to characterize dendritic spine morphology in pyramidal neurons from the ACC. Rodents with a history of adolescent OXY self-administration exhibited a sex-dependent population shift in the morphological parameters of dendritic spines (Backbone Length: OXY Access x Sex x Bin Interaction, [*F*(1,699)=295.8, *p*≤0.001]; Head Diameter: OXY Access x Sex x Bin Interaction, [*F*(1,403)=289.4, *p*≤0.001]). Complementary analyses were conducted by sex to establish the locus of the interaction.

Male rats with a history of adolescent OXY self-administration (Figure 4) exhibited a population shift towards dendritic spines with increased backbone length (Figure 4A; OXY Access x Bin Interaction, [*F*(1,397)=180.5, *p*≤0.001]) and decreased head diameter (Figure 4B; OXY Access x Bin Interaction, [*F*(1,229)=90.4, *p*≤0.001]) relative to control animals. Regression analyses further revealed the dose-dependency of dendritic spine dysmorphology in male animals, whereby a male rodent’s average OXY intake during the non-abstinent period prior to euthanasia was linearly related to median backbone length (Figure 4C; *R*^2^=0.375, Null Hypothesis, β_1_=0: [*F*(1,19)=11.4, *p*≤0.001]) and median head diameter (Figure 4D; *R*^2^=0.220, Null Hypothesis, β_1_=0: [*F*(1,19)=5.4, *p*≤0.032]). As a result, 37.5% and 22.0% of the variance in median backbone length and head diameter, respectively, was accounted for by a male rodent’s average OXY intake during the non-abstinent period prior to euthanasia.

**Figure 4.**
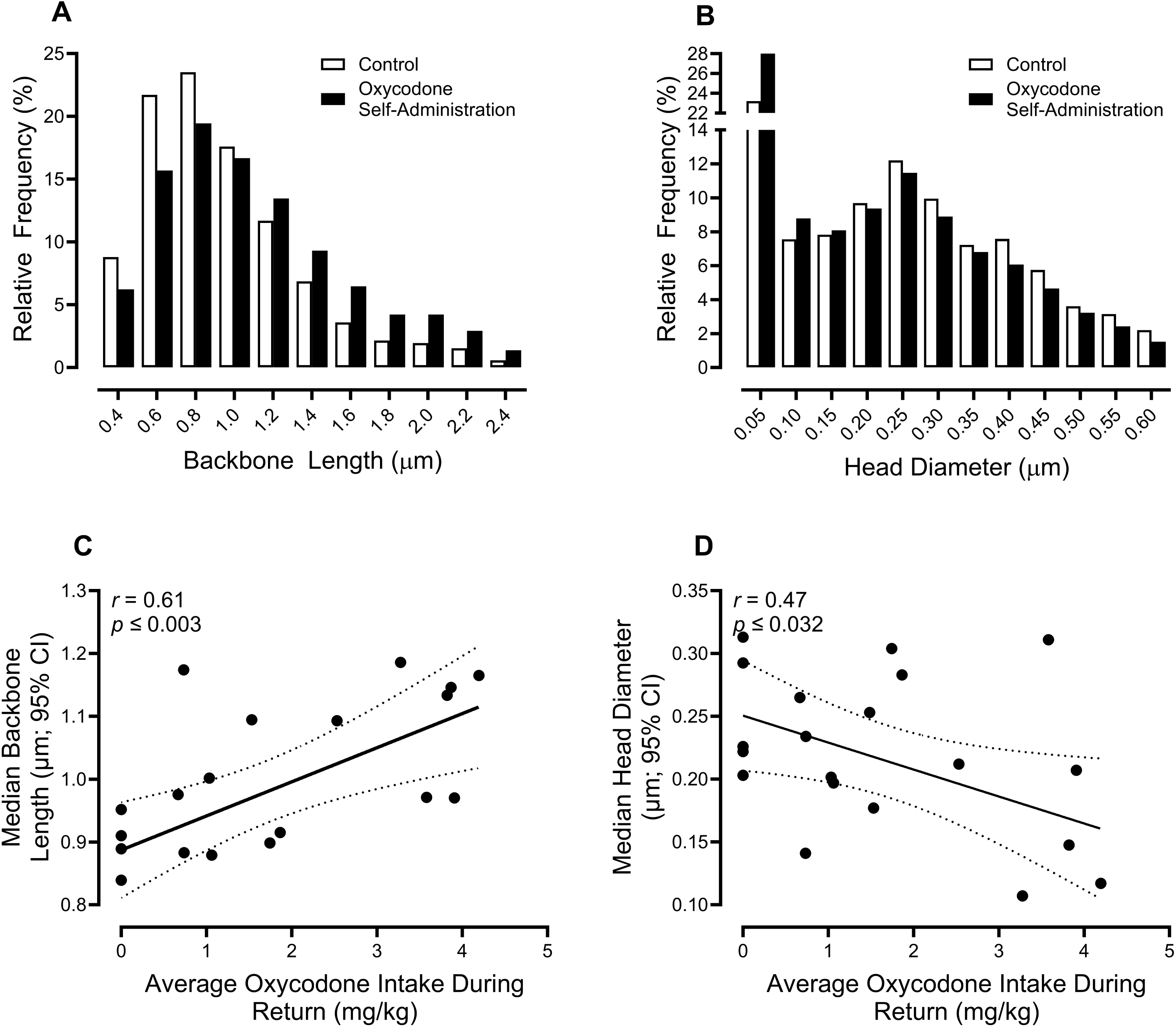
Male Dendritic Spine Morphology. (**A,B**) In male animals, a history of adolescent oxycodone (OXY) self-administration induced a statistically significant (*p* ≤ 0.05) population shift towards increased dendritic spine backbone length and decreased dendritic spine head diameter (Control (i.e., 0 mg/kg): *n*=5; Oxycodone Self-Administration: *n*=16 (2 mg/kg: *n*=5, 5 mg/kg: *n*=5, 10 mg/kg: *n*=6)). Data are presented as relative frequencies (%). (**C**,**D**) Alterations in dendritic spine morphology in male rodents are dose-dependent, whereby a male rodent’s average OXY intake during the non-abstinent period prior to euthanasia was positively associated with median backbone length (*r* = 0.61, *R*^2^=0.37, *p* ≤ 0.003) and negatively related to median head diameter (*r* = 0.47, *R*^2^=0.22, *p* ≤ 0.032). 95% Confidence Intervals for the regression are illustrated with dotted lines.

In sharp contrast, a population shift towards dendritic spines with decreased backbone length (Figure 5A; OXY Access x Bin Interaction, [*F*(1,302)=117.0, *p*≤0.001]) and increased head diameter (Figure 5B; OXY Access x Bin Interaction, [*F*(1,403)=289.4, *p*≤0.001]) was observed in female rodents with a history of adolescent OXY self-administration relative to control rats. Alterations in dendritic spine morphology in female animals were dose-dependent, whereby a female rodent’s average OXY intake during adolescence was linearly associated with the 75^th^ percentile of backbone length (Figure 5C; *R*^2^=0.256, Null Hypothesis, β_1_=0: [*F*(1,14)=4.8, *p*≤0.05]) and head diameter (Figure 5D; *R*^2^=0.332, Null Hypothesis, β_1_=0: [*F*(1,14)=6.9, *p*≤0.02]). Accordingly, a female rodent’s average OXY intake during adolescence accounted for 25.6% of the variance in the 75^th^ percentile of backbone length and 33.2% of the variance in the 75^th^ percentile of head diameter. Taken together, OXY self-administration sex-and dose-dependently altered dendritic spine morphology, with higher doses of self-administration inducing greater adverse effects.

**Figure 5.**
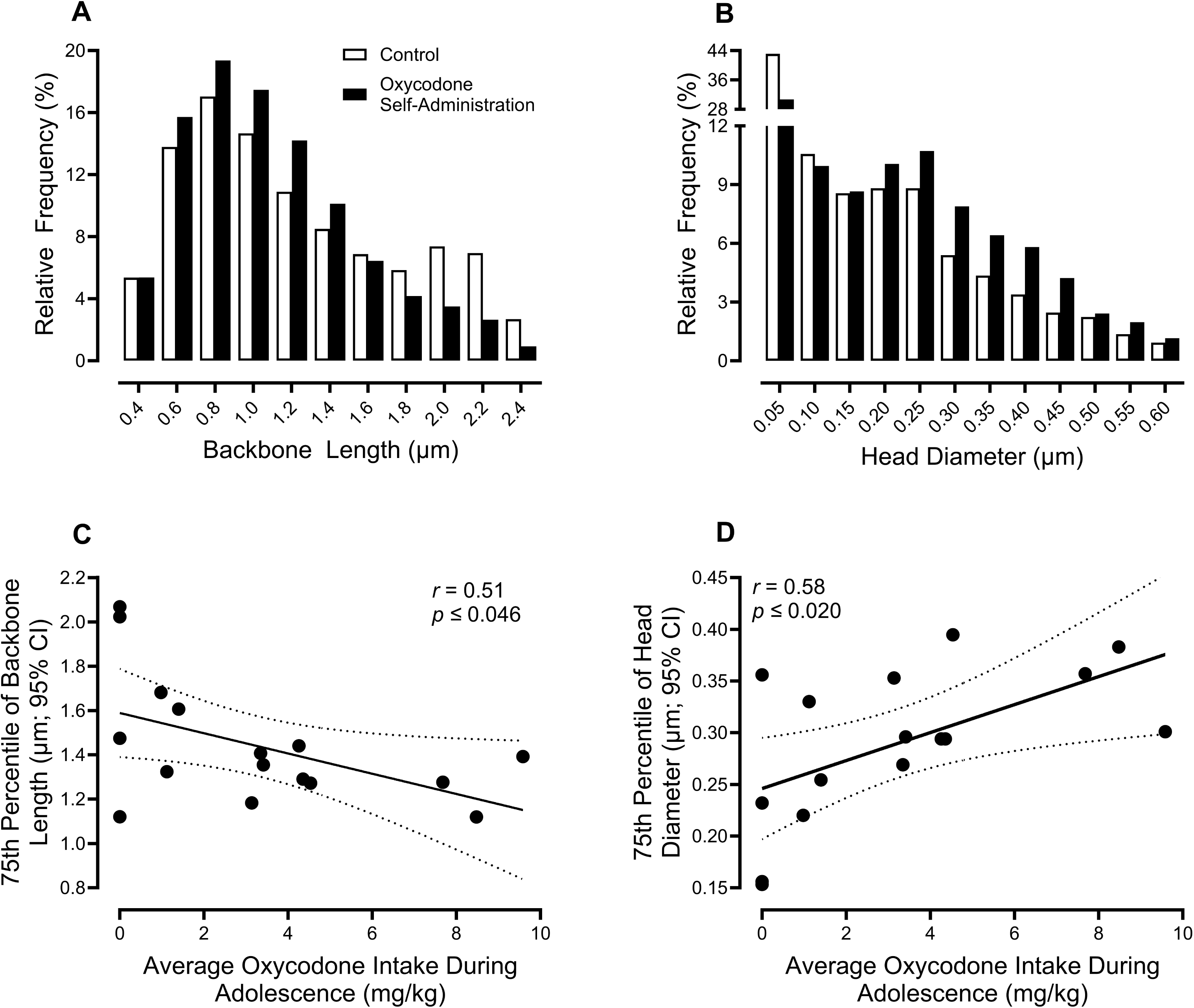
Female Dendritic Spine Morphology. (**A,B**) In female rodents, adolescent oxycodone (OXY) self-administration induced a statistically significant (*p* ≤ 0.05) population shift towards decreased dendritic spine backbone length and increased dendritic spine head diameter (Control (i.e., 0 mg/kg): *n*=4; OXY Self-Administration: *n*=12 (2 mg/kg: *n*=3; 5 mg/kg: *n*=4; 10 mg/kg: *n*=5)). Data are presented as relative frequencies (%). (**C,D**) Dendritic spine morphological alterations in female rats are dose-dependent, whereby a female animal’s average OXY intake during adolescence was negatively and positive related to the 75^th^ Percentile of Backbone Length (*r*=0.51, *R*^2^=0.26, *p* ≤ 0.046) and Head Diameter (*r*=0.58, *R*^2^=0.34, *p* ≤ 0.020), respectively. 95% Confidence Intervals for the regression are illustrated with dotted lines.

### Rodents with a history of adolescent oxycodone self-administration exhibited selective, sex-dependent alterations in genes encoding synaptic proteins and glutamate receptors in the prefrontal cortex

Transcriptional changes in the mRNA expression of 12 genes in the PFC were evaluated as a proxy for synaptic scaffolding proteins (i.e., *dlg3, dlg4*), a protein kinase localized to postsynaptic compartments (i.e., *cdk5*^32–33^), NMDARs (i.e., *GRIN1, GRIN2A, GRIN2C, GRIN2D, GRIN3B*), and AMPARs (i.e., *GRIA1, GRIA2, GRIA3, GRIA4*). mRNA expression was selectively altered by OXY self-administration (Figure 6A-D; Supplemental Figure 1).

**Figure 6.**
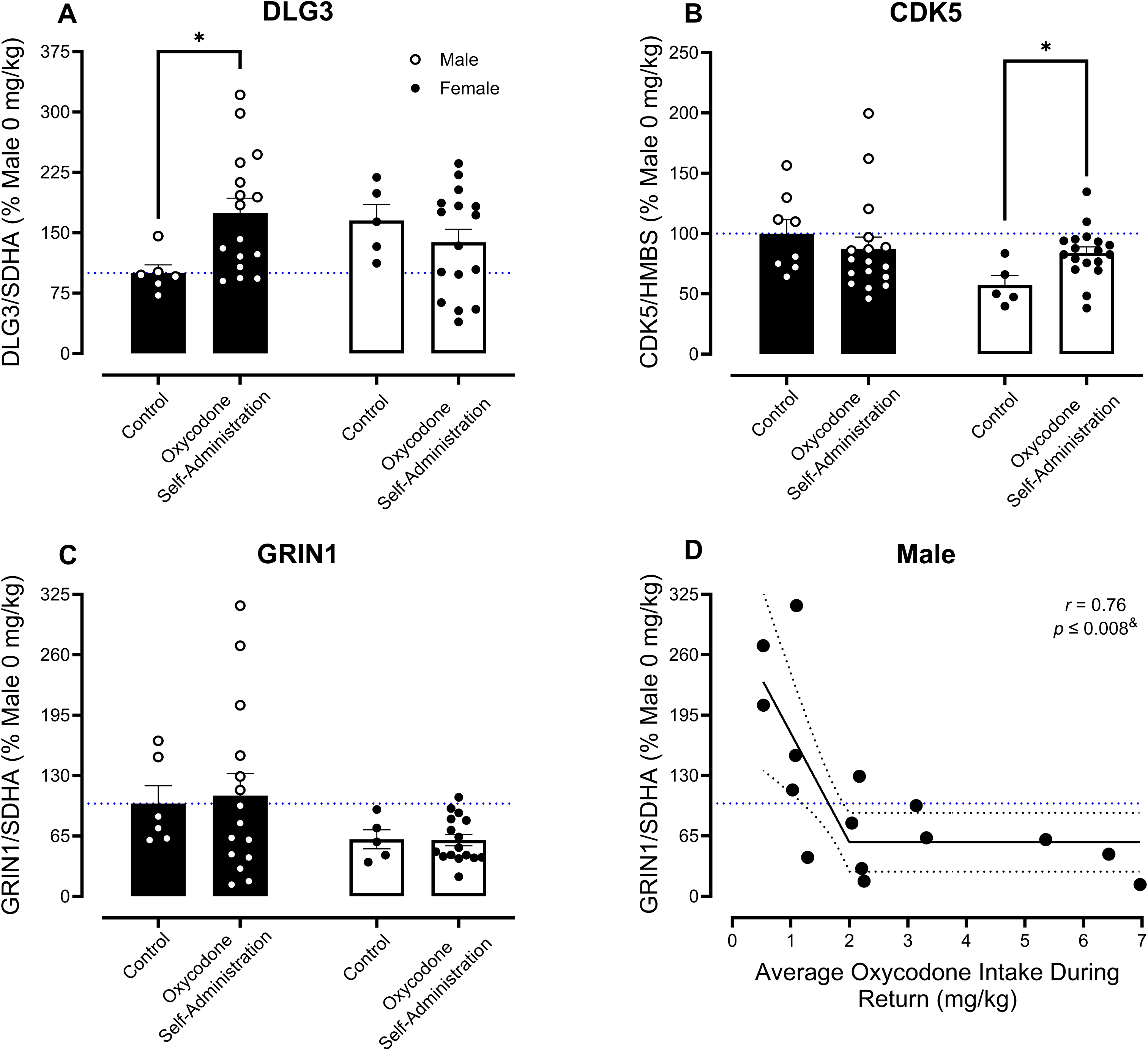
Quantitative Polymerase Chain Reaction (qPCR). (**A,B,C**) The bar graphs (Mean (X) ± Standard Error of the Mean (SEM)) illustrate qPCR analysis of *Dlg3, Cdk5,* and *GRIN1* mRNA expression in the prefrontal cortex. A history of adolescent oxycodone (OXY) self-administration significantly increased *Dlg3* and *Cdk5* mRNA expression in male and female rodents, respectively. *GRIN1* mRNA expression is not significantly altered by adolescent OXY self-administration in male or female animals. Sample sizes for qPCR analyses are as follows: Control (i.e., 0 mg/kg), *n*=11-13 (male, *n*=6-8, female, *n*=5), OXY Self-Administration, *n*=32-35 (male, *n*=16-17, female, *n*=16-18) (**D**) Nevertheless, *GRIN1* mRNA expression is dose-dependently altered in male rodents, whereby the relationship between *GRIN1* mRNA expression and a male rodent’s average OXY intake during the non-abstinent period prior to euthanasia is well-described by a segmental linear regression (*r*=0.76, *R*^2^=0.58, *p* ≤ 0.008). Confidence Intervals for the regression are illustrated with black dotted lines. The blue dotted line represents 100% mRNA expression for control male animals. \**p* ≤ 0.05.

*Dlg3*, a scaffold protein involved in the structure and function of excitatory synapses, and *cdk5,* a kinase involved in pre- and postsynaptic function, were altered in a sex-dependent manner by a history of adolescent oral OXY self-administration (Figure 6A-B; *dlg3:* OXY Access x Sex Interaction, [*F*(1,39)=5.9, *p*≤0.02, η_p_^2^=0.131]; *cdk5*: OXY Access x Sex Interaction, [*F*(1,44)=6.4, *p*≤0.015, η_p_^2^=0.126]). Complementary analyses were conducted by sex to elucidate the locus of the interaction. A prominent decrease in *dlg3* expression in the PFC was observed in male (Main Effect: OXY Access, [*F*(1,20)=7.3, *p*≤0.014, η_p_^2^=0.268]), but not female (*p*>0.05), OXY rodents relative to male or female control animals, respectively. With regards to *cdk5* expression, female (Main Effect: OXY Access, [*F*(1,21)=7.0, *p*≤0.015, η_p_^2^=0.250]), but not male (*p*>0.05), OXY animals exhibited a profound increase in *cdk5* expression relative to female or male control animals, respectively. Alterations in neither *dlg3* or *cdk5* expression in the PFC were not, however, dose-dependent (Data Not Shown; OXY Dose x Sex Interaction, *p*>0.05).

Adolescent OXY self-administration sex- and dose-dependently altered mRNA expression of *GRIN1*, a gene that encodes for a subunit of the NMDA receptor. Presence or absence of OXY self-administration during adolescence failed to alter mRNA expression of *GRIN1* (Figure 6C; OXY Access x Sex Interaction, *p*>0.05; Main Effect: OXY Access, *p*>0.05). *GRIN1* expression, however, was sex- and dose-dependently changed by a history of adolescent OXY self-administration (OXY Dose x Sex Interaction, [*F*(3,34)=5.1, *p*≤0.005, η_p_^2^=0.312]). Complementary analyses, which were conducted by sex, revealed the locus of the interaction, whereby male (Main Effect: OXY Dose, [*F*(1,17)=5.4, *p*≤0.009, η_p_^2^=0.487]), but not female (*p*>0.05), OXY rodents exhibited a prominent dose-dependent decrease in *GRIN1* expression relative to male or female control animals, respectively. A segmental linear regression constrained at 2 mg/kg (Figure 6D; Null Hypothesis, Slope 1=0: [*F*(1,12)=10.0, *p*≤0.01]; Null Hypothesis, Slope 2=0: [*F*(1,12)=1.6, *p*>0.05]) confirmed these observations, with a male rodent’s average OXY intake during the non-abstinent period prior to euthanasia accounting for 57.7% of the variance in *GRIN1* expression.

A history of adolescent oral OXY self-administration failed to alter mRNA expression of *dlg4, GRIN2A, GRIN2C, GRIN2D, GRIN3B, GRIA1, GRIA2, GRIA3, or GRIA4* (Supplemental Figure 1; OXY Access x Sex Interaction, *p*>0.05; Main Effect: OXY Access, *p*>0.05; OXY Dose x Sex Interaction, *p*>0.05; Main Effect: OXY Dose, *p*>0.05). Notably, however, the factor of biological sex influenced mRNA expression of *GRIN2D* (Main Effect: Sex, [*F*(1,38)=11.2, *p*≤0.002, η_p_^2^=0.227]), *GRIA1* (Main Effect: Sex, [*F*(1,43)=5.6, *p*≤0.022, η_p_^2^=0.116]), and *GRIA3* (Main Effect: Sex, [*F*(1,38)=4.1, *p*≤0.05, η_p_^2^=0.098]).

Collectively, male rodents with a history of OXY self-administration exhibited increased *dlg3* expression and a dose-dependent decrease in *GRIN1* expression. In sharp contrast, a significant increase in *cdk5* expression was observed in female animals with a history of OXY self-administration. Results support, therefore, selective, sex-dependent alterations in the GluN1 subunit of the NMDA receptor and *cdk5*.

### Alterations in mRNA expression and dendritic spine dysmorphology partially underlie cognitive impairments in animals with a history of adolescent prescription opioid use disorder

Regression analyses were subsequently conducted to evaluate if the alterations in transcriptomic expression and/or dendritic spine dysmorphology mechanistically underlie previously published APOUD-induced impairments in sensorimotor gating, indexed using visual PPI, or locomotion, indexed using open field activity.^11^

NMDA receptor activity is fundamentally involved in the regulation of dendritic spine morphology, whereby *dlg3* expression promotes the elongation of dendritic spines in an NMDA receptor activity-dependent manner.^44^ Indeed, consistent with prior studies,^44^ *GRIN1* expression is linearly related to *dlg3* expression in male rodents, with *GRIN1* expression accounting for 33.6% of the variance in *dlg3* expression (Figure 7A; [*F*(1,12)=6.1, *p*≤0.03]). The fundamental role of dendritic spine dysmorphology in OXY-induced neurocognitive impairments was subsequently evaluated, whereby *GRIN1* expression (Figure 7B; [*F*(1,17)=5.6, *p*≤0.03]) and structural alterations in dendritic spines (i.e., Median Backbone Length, Median Head Diameter, Figure 7C; [*F*(2,17)=4.6, *p*≤0.03]) accounted for 34.5% and 35.0% of the variance, respectively, in visual PPI at PD 180. In male animals, therefore, OXY-induced alterations in NMDA receptor activity likely induce a population shift towards longer dendritic spine backbone length, resulting in the development of neurocognitive impairments.

**Figure 7.**
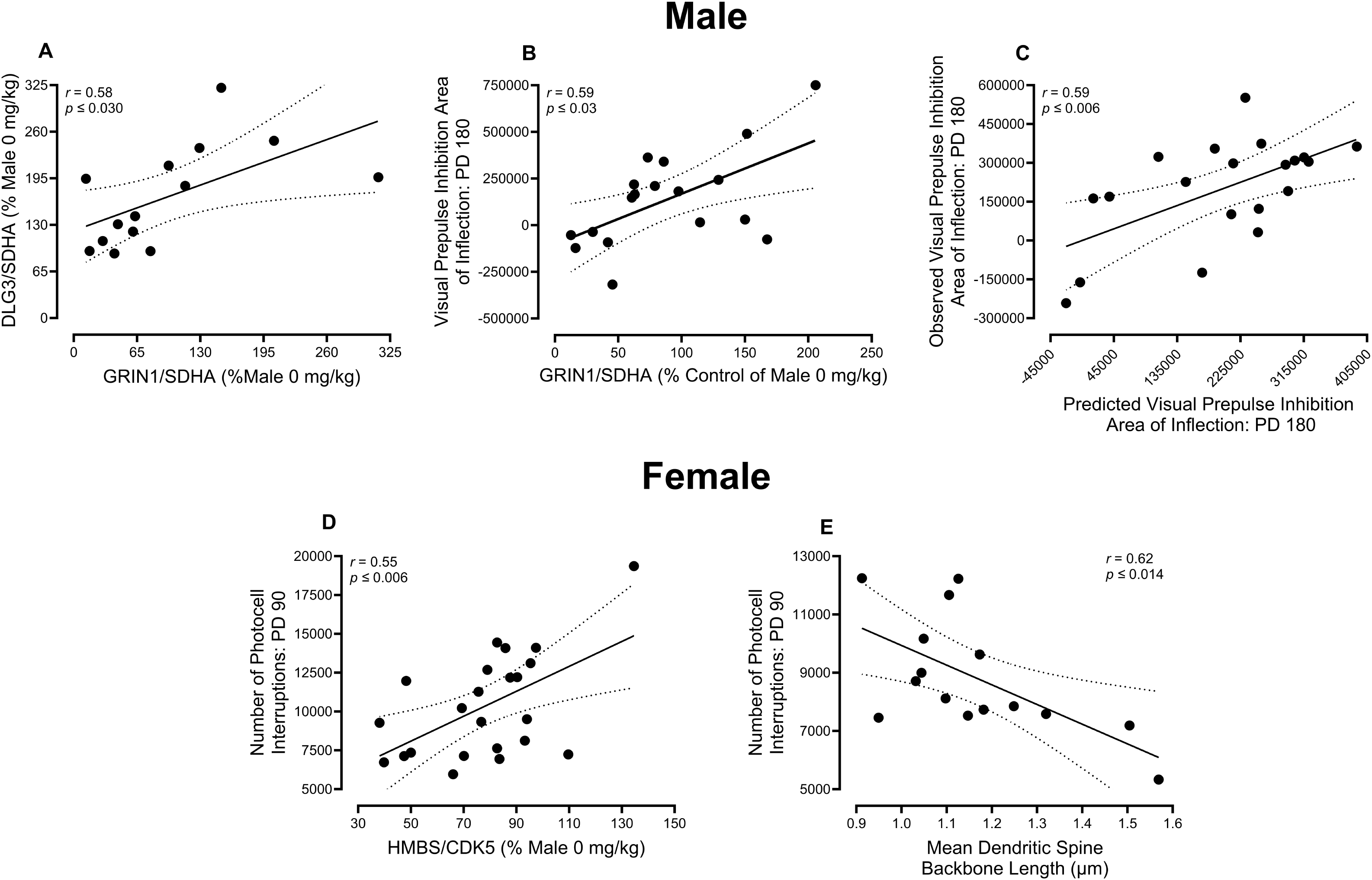
Regression Analysis. (**A**) In male rodents, a linear regression analysis was conducted to evaluate the relationship between *Dlg3* and *GRIN1* mRNA expression. Increased *GRIN1* mRNA expression was significantly associated with increased *dlg3* mRNA expression (*r*=0.58, *R*^2^=0.34, *p* ≤ 0.030). (**B,C**) Linear and multiple regression analyses in male animals were conducted to evaluate if and/or how *Dlg3* and dendritic spine dysmorphology were related to neurocognitive impairments in preattentive processes at postnatal day (PD) 180 (previously reported by McLaurin et al. [11]). Indeed, *Dlg3* mRNA expression and structural alterations in dendritic spines (i.e., Median Backbone Length, Median Head Diameter) accounted for 50.4% (*r*=0.71, *p* ≤ 0.003) and 35.0% (*r*=0.59, *p* ≤ 0.006) of the variance in visual prepulse inhibition. (**D**) In female rats, linear regression analyses were conducted to evaluate if and/or how *Cdk5* and dendritic spine dysmorphology were related to neurocognitive impairments in locomotion at PD 90. Indeed, *Cdk5* mRNA expression and mean dendritic spine backbone length accounted for 30.2% (*r*=0.55, *p* ≤ 0.006) and 38.4% (*r*=0.62, *p* ≤ 0.014) of the variance in open field activity at PD 90.

In female rodents, *cdk5* expression and structural alterations in dendritic spines mechanistically underlie the prominent neurocognitive impairments exhibited by female rats with a history of oral adolescent OXY self-administration. Indeed, independently, *cdk5* expression (Figure 7D; [*F*(1,21)=9.2, *p*≤0.006] and the mean dendritic spine backbone length (Figure 7E; *F*(1,13)=8.1, *p*≤0.014) accounted for 30.1% and 38.4% of the variance in open field activity at PD 90, respectively.

Taken together, mRNA expression and structural alterations in dendritic spines in pyramidal neurons from the ACC afford a sex-dependent neural mechanism underlying neurocognitive impairments induced by APOUD.

## DISCUSSION

Chronic oral adolescent OXY self-administration induces profound sex-dependent impairments in dendritic spine dysmorphology and transcriptional dysregulation in the PFC. A ballistic labeling technique, in concert with sophisticated neuronal reconstruction software, was utilized to measure the morphological parameters (i.e., backbone length, head diameter) of over 54,000 dendritic spines located on pyramidal neurons in the ACC. A prominent sex- and dose-dependent population shift in the morphological parameters (i.e., backbone length, head diameter) of dendritic spines was induced by a history of oral adolescent OXY self-administration, supporting profound synaptic dysfunction. qPCR was subsequently utilized to investigate the mechanism underlying dendritic spine dysmorphology. Transcriptional changes in the mRNA expression of *GRIN1,* a gene that encodes for the GluN1 subunit of the NMDA receptor, were observed in male, but not female, rodents with a history of oral adolescent OXY self-administration relative to their control counterparts. In sharp contrast, in female rats, a history of oral adolescent OXY self-administration induced transcriptional alterations in the mRNA expression of *Cdk5*, a protein kinase that responds to excessive glutamate concentrations.^45^ Fundamentally, regression analyses revealed a strong association (i.e., *r* = 0.55 – 0.71) between either biochemical dysregulation or dendritic spine dysmorphology and neurocognitive processes; an association that supports key neural mechanisms underlying APOUD-induced neurocognitive impairments. Taken together, results support that sex-dependent glutamatergic dysregulation may underlie dendritic spine dysmorphology in the ACC resulting in APOUD-induced neurocognitive impairments. Targeting glutamatergic dysregulation via novel therapeutics may mitigate neurocognitive impairments resulting from a history of adolescent OXY self-administration.

Elucidating alterations in dendritic spine morphology affords key insights into the ongoing structural and functional adaptations that occur consequent to chronic oral APOUD. Dendritic spine nomenclature traditionally classifies spines into one of four fixed categories (i.e., filopodia, mushroom, stubby, or thin) based on their morphological characteristics.^46^ Filopodia and thin dendritic spines are highly transient, long (i.e., increased backbone length) protrusions that lack a distinct spine head or have a small head, respectively. Mushroom spines, which are associated with strong, long-lasting synapses, are characterized by a large, bulbous dendritic spine head and a thin dendritic spine neck. Stubby dendritic spines, in sharp contrast, have a wide spine head, but are devoid of a dendritic spine neck.^46–47^ Despite the inherent simplicity in dendritic spine classification, the advantages of exploring dendritic spine morphology along a continuum^48^ cannot be understated, as it affords a fundamental opportunity to infer synaptic function. Indeed, dendritic spine backbone length is inversely associated with excitatory postsynaptic potentials, an index of synaptic efficacy,^49^ whereas the size of the dendritic spine head is positively correlated with area of the PSD.^28–29^ Thus, the prominent sex- and dose-dependent alteration in dendritic spine morphology observed in rodents with a history of chronic oral adolescent OXY self-administration supports profound impairments in synaptic function in pyramidal neurons in the ACC.

The potential role of synaptodendritic dysfunction in the pathophysiology of opioid use disorders, and associated consequences (e.g., neurocognitive impairments), cannot be understated, as both naturally derived opiates (e.g., morphine)^50–51^ and synthetic opioids (e.g., OXY; ^52^, ^Present^ ^Study^ Fentanyl)^53^ induce sex- and dose-dependent changes in neuronal morphology, dendritic spine density, and dendritic spine morphology. Notably, however, a number of factors, including delivery mechanism (i.e., experimenter delivered^50–51^ versus self-administration)^50^ and mu opioid receptor (MOR) internalization^54^ influence precisely how neuronal and/or dendritic spine dysmorphology is expressed. For example, experimenter delivered morphine decreases neuronal arbor complexity and dendritic spine length in adult male rodents^51^; findings that are in contrast to those observed in male rodents with a history of oral adolescent OXY self-administration. Nevertheless, herein, findings of treatment- and sex-dependent reductions in neuronal arbor complexity replicate those previously reported in striatal neurons exposed to OXY *in vitro*.^52^ Furthermore, opioids that have a weak or non-internalizing effect on the MOR (i.e., Morphine, OXY, Low Concentrations of Fentanyl) consistently decrease dendritic spine density,^50,52–53^ whereas strong MOR internalization (i.e., High Concentrations of Fentanyl) increases dendritic spine density.^53^ Despite the valuable information garnered from preclinical studies of a single substance, translational studies examining dendritic spine density in humans with ongoing opioid use disorder are fundamentally needed; the recent development of novel tools to measure *in vivo* synaptic density (e.g., Synaptic Vesicle Protein 2A Positron Emission Tomographic Imaging) provides an approach to address this, as of yet, unmet need.

Consistent with the profound sex-dependent dendritic spine dysmorphology induced by a history of chronic oral adolescent OXY self-administration, the biochemical mechanism underlying these alterations is dependent upon the factor of biological sex. NMDARs, which are postsynaptic glutamate receptors, are heterotetramers assembled from the GluN1 (i.e., encoded by the *GRIN1* gene) subunit, four distinct GluN2 subunits (i.e., 2A, 2B, 2C, and 2D), and two GluN3 subunits (i.e., 3A and 3B). Selective structural alterations in NMDARs were evidenced by a dose-dependent decrease in *GRIN1* mRNA expression in male rats with a history of oral adolescent OXY self-administration. The GluN1 subunit of male rodents appears uniquely vulnerable to insult during adolescence, as both adolescent isolation rearing^55^ and adolescent exposure to delta-9-tetrahydrocannabinol^56^ also decrease expression of NMDA GLuN1 subunits. From a functional perspective, the activity of NMDARs regulates dendritic spine morphology via ionotropic^57^ and non-ionotropic^58–59^ signaling pathways, as well as scaffolding proteins (e.g., SAP102, PSD-95).^44^ Indeed, alternative splicing of the scaffolding protein SAP102, which is encoded by the *dlg3* gene, directly regulates dendritic spine morphology, whereby the N-terminal splice variant promotes lengthening of dendritic spines,^44^ as observed in male animals with a history of chronic oral adolescent OXY self-administration.

The prominent neuronal and dendritic spine dysmorphology observed in female rodents with a history of adolescent OXY self-administration, however, may result from hyperactivity of *cdk5*. *Cdk5* is a predominantly neural-specific serine/threonine kinase activated by p35 and p39.^60–61^ Activity of the neuronal *cdk5* pathway is tightly regulated, whereby either inactivation or hyperactivation of *cdk5* dysregulates the neuronal cytoskeleton.^62^ Indeed, inhibition of *cdk5* activity hinders neurite outgrowth^63–65^ and impairs the proper maturation of dendritic spines.^63^ To date, however, no studies have systematically investigated how *cdk5* hyperactivation influences dendritic spine morphology. Nevertheless, high levels of glutamate, which have been observed in clinical subjects with POUD,^66–67^ trigger the activation of *cdk5*,^45^ inhibits dendrite growth,^68–70^ and rapidly alters dendritic spine morphology.^71^ More specifically, with regards to dendritic spines, high levels of glutamate lead to the loss of filamentous actin^72^ resulting in an increased proportion of stubby spines;^73^ findings that resemble the OXY-induced dendritic spine dysmorphology observed in female rodents. Transcription of *cdk5* appears vulnerable to opioid exposure, whereby both inactivation^74–75^ and hyperactivation^74^ have been reported. Although experimental protocol limitations preclude direct associations between mRNA expression and dendritic spine morphology, findings of the present study in concert with prior work support alterations in *GRIN1* or *cdk5* as a mechanism partly underlying the neuronal and dendritic spine dysmorphology in male and female rodents, respectively, following adolescent OXY self-administration.

Fundamentally, sex-dependent biochemical dysregulation and dendritic spine dysmorphology partly underlie APOUD-induced neurocognitive impairments, evidenced by simple linear and multiple regression analyses. Indeed, *GRIN1* expression and the morphological parameters of dendritic spines (i.e., backbone length, head diameter) in male rodents accounted for 34.5% and 35.0%, respectively of the variance in visual PPI at PD 180. It is notable, albeit unsurprising, that dendritic spine dysmorphology in male animals would be associated with the final neurocognitive evaluation of visual PPI, as recent learning and memory formation are reflected in the structural parameters of dendritic spines. Consistently, a male rodent’s average OXY intake during the non-abstinent period immediately prior to euthanasia was also associated with dendritic spine backbone length and head diameter. More surprisingly, however, in female rats, *cdk5* expression and the mean dendritic spine backbone length accounted for 30.1% and 38.4%, respectively of the variance in open field activity at PD 90, a neurocognitive assessment conducted following chronic OXY self-administration, but prior to spontaneous withdrawal. In female rats, average OXY intake during adolescence was also significantly associated with dendritic spine morphology. Biochemical dysregulation and dendritic spine dysmorphology in female rodents, therefore, likely reflects long-lasting neuroadaptations induced by chronic adolescent oral OXY self-administration, whereas alterations in male rats may be due to oral OXY self-administration during the non-abstinence period.

Taken together, the present study provides fundamental insight into the neural mechanisms underlying APOUD-induced dendritic spine dysmorphology and neurocognitive impairments. Indeed, dendritic spine dysmorphology in male and female rodents with a history of oral adolescent OXY self-administration is characterized by a population shift towards a more immature dendritic spine phenotype (i.e., thin and stubby dendritic spines, respectively). Results of the present study, in concert with prior literature, support transcriptional dysregulation of glutamatergic associated genes (i.e., *GRIN1, cdk5*) as a biochemical mechanism partly underlying dendritic spine dysmorphology. Fundamentally, both transcriptional dysregulation and dendritic spine dysmorphology are associated with measures of APOUD-induced neurocognitive impairments. Thus, targeting glutamatergic dysregulation via novel therapeutics may mitigate neurocognitive impairments resulting from a history of adolescent OXY self-administration.

## AUTHOR CONTRIBUTIONS

**Kristen A. McLaurin:** Conceptualization, Funding acquisition, Data curation, Writing-original draft, Writing-reviewing & editing, Formal analysis. **Taylor Elder:** Data curation, Writing-reviewing & editing. **Hailong Li:** Data curation, Writing-reviewing & editing. **Brendan J. Veenstra:** Data curation, Writing-reviewing & editing. **Kyler J. Nelson:** Data curation, Writing-reviewing & editing. **Charles F. Mactutus:** Writing-reviewing & editing, Funding acquisition, Conceptualization. **Jill R. Turner:** Writing-reviewing & editing, Funding acquisition. **Rosemarie M. Booze:** Writing-reviewing & editing, Funding acquisition, Conceptualization.

## ACKNOWLEDGEMENTS

No author has an actual or perceived conflict of interest with the contents of this article.

## DATA AVAILABILITY STATEMENT

The authors declare that all the data supporting the findings of this study are available within the paper and its Supplemental Data.

## Non-Standard Abbreviation List

ACC: Anterior Cingulate Cortex
AMPAR: α-Amino-3-hydroxy-5-methyl-4-isoxazolepropionic acid receptors
ANOVA: Analysis of Variance
APOUD: Adolescent Prescription Opioid Use Disorder
CDK5: Cyclin-Dependent Kinase 5
HMBS: Hydroxymethylbilane Synthase
MOR: Mu Opioid Receptor
mPFC: Medial Prefrontal Cortex
NMDAR: N-methyl-D-aspartate receptors
PBS: Phosphate Buffered Saline
PD: Postnatal Day
PFC: Prefrontal Cortex
POUD: Prescription Opioid Use Disorder
PPI: Prepulse Inhibition
PSD: Postsynaptic Density
qPCR: Quantitative Polymerase Chain Reaction
SDHA: Succinate Dehydrogenase Complex Subunit A

