## Supplementary Information for "A HISTORY OF ADOLESCENT OXYCODONE SELF-ADMINISTRATION: NEURAL MECHANISMS UNDERLYING NEUROCOGNITIVE IMPAIRMENTS"

789 South Limestone Street

Lexington, KY 40508

^2^Cognitive and Neural Science Program

Department of Psychology

Barnwell College

University of South Carolina

1512 Pendleton Street

Columbia, SC 29208

**Number of Tables, Figures**: 1, 6

**Running Head (≤48 characters):** Oxycodone Induced Dendritic Spine Dysmorphology

**Address proofs and correspondence to:**

Kristen A. McLaurin, Ph.D.

Assistant Professor

Department of Pharmaceutical Sciences

College of Pharmacy

789 South Limestone Street

University of Kentucky

Lexington, KY 40508

**Non-Standard Abbreviation List:**

ACC: Anterior Cingulate Cortex

AMPAR: α-Amino-3-hydroxy-5-methyl-4-isoxazolepropionic acid receptors

ANOVA: Analysis of Variance

APOUD: Adolescent Prescription Opioid Use Disorder

CDK5: Cyclin-Dependent Kinase 5

HMBS: Hydroxymethylbilane Synthase

MOR: Mu Opioid Receptor

mPFC: Medial Prefrontal Cortex

NMDAR: N-methyl-D-aspartate receptors

PBS: Phosphate Buffered Saline

PD: Postnatal Day

PFC: Prefrontal Cortex

POUD: Prescription Opioid Use Disorder

PPI: Prepulse Inhibition

PSD: Postsynaptic Density

qPCR: Quantitative Polymerase Chain Reaction

SDHA: Succinate Dehydrogenase Complex Subunit A

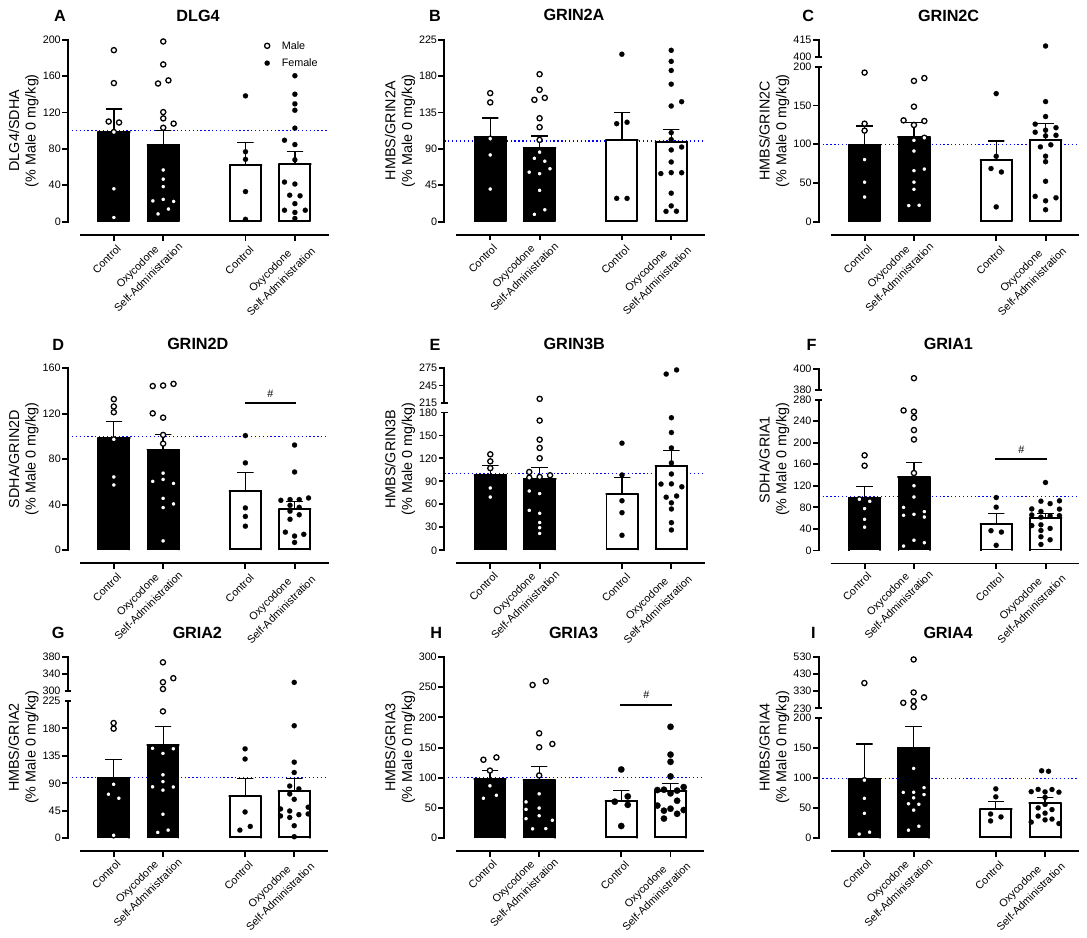

**Supplemental Figure 1.** Quantitative Polymerase Chain Reaction (qPCR). (**A-I**) The bar graphs (Mean (X) ± Standard Error of the Mean (SEM)) illustrate qPCR analysis of proxies for synaptic function (i.e., *dlg4*), NMDARs (i.e., *GRIN2A, GRIN2C, GRIN2D, GRIN3B*), and AMPARs (i.e., *GRIA1, GRIA2, GRIA3, GRIA4*). A history of adolescent oral oxycodone self-administration failed to alter mRNA expression. The factor of biological sex, however, influenced mRNA expression of *GRIN2D*, *GRIA1,* and *GRIA3*. The blue dotted line represents 100% mRNA expression for control male animals.
